# Rational Design of Molecular Glue Degraders Utilizing Endogenous Transient Protein–Protein Interactions

**DOI:** 10.64898/2026.11.16.766899

**Authors:** Zilu Li, Suyu Ji, Mi Ke, Wenyu Zhu, Xiaolei Zhang, Yipu Chen, Yingfan Liu, Peng Zhang, Yuying Ye, Gaoang Wang, Jiahui Zhu, Yiting Chu, Xinyu Sun, Wensi Zhao, Minjia Tan, Mingliang Ye, Yanyan Lan, Ruijun Tian, Ke Ding

**Author notes:** These authors contributed equally: Zilu Li, Suyu Ji, Mi Ke, Wenyu Zhu, Xiaolei Zhang.

## Abstract

Molecular glues are small molecules that modulate protein activity by inducing or stabilizing protein–protein interactions (PPIs). However, identifying and harnessing endogenous protein pairs amenable to small-molecule-mediated stabilization remains a major challenge in molecular glue development. Here we introduce transient interface pocket-guided molecular glue rational design (TrIP–MGRD), a strategy that shifts compound discovery from empirical screening toward rational, interaction-guided design by exploiting druggable pockets at transient PPI interfaces. Using CDK12 as a model target, we identified DDB1 as an endogenous transient interaction partner and determined the DDB1–CDK12–CCNK complex structure by cryo-electron microscopy (cryo-EM). Structure-guided virtual screening and medicinal chemistry optimization yielded a molecular glue C432 that stabilized this interaction. Comparison of the ligand-free and C432-bound complexes revealed pre-existing contacts at the transient CDK12–DDB1 interface provides a structural basis for compound-mediated stabilization. Collectively, by integrating endogenous partner identification with interface-guided screening, TrIP–MGRD provides a new framework for target-directed molecular glue discovery.

## Introduction

PPIs underpin cellular organization and regulate a wide range of biological processes^1^. Based on their binding patterns, PPIs are broadly classified as permanent or transient^2^. Transient PPIs are widespread yet incompletely characterized. Unlike permanent PPIs, transient protein–protein partner recruitment was controlled by conformational changes, post-translational modifications exposing or remodeling protein-binding surfaces, which modulates the biological functions^3,4^. Recent biochemical and biophysical studies have shown that in some molecular glue degrader systems, the E3 ligase and protein of interest (POI) can interact transiently in the absence of the compound. Molecular-glue binding can reshape protein recognition surfaces and strengthen these interactions, thereby stabilizing ternary complexes^5-7^. These findings highlight the potential of harnessing transient PPIs to modulate cellular function.

Unlike conventional inhibitor screening, molecular-glue discovery requires identifying both a compatible protein pair and a ligand that stabilizes their interaction. The main strategies for discovering molecular glues include identifying compound-induced interactions through phenotypic, proteomic or affinity-based screening, which yielded the GSPT1 degrader CC-885^8^, CRBN-based KAT2A degraders^5^ and the DDX18 degrader M12^9^. FKBP12-focused libraries have also yielded BRD9 and QDPR recruiters^10^. Some strategies screen ligands against biologically established protein pairs, exemplified by fluorescence polarization screening of β-TrCP–β-catenin enhancers subsequently optimized into NRX-252114^11^ and disulfide fragment tethering of 14-3-3σ–ERα^12^. Despite these advances, a broadly applicable strategy to identify suitable endogenous transient PPI pairs before compound screening, and to assess whether their interfaces contain pockets suitable for molecular-glue binding is still lacking.

Here, we present TrIP–MGRD, a workflow that exploits suitable endogenous transient PPIs as starting points for molecular-glue discovery. Using H_2_O_2_-independent ROProx proximity labeling in living cells^13^, we identified candidate endogenous partners DDB1 of CDK12. Combined with peptide-centric local stability assay (PELSA)^14^, crosslinking-assisted cryo-EM enabled structural characterization of the transient CDK12–DDB1 complex, providing an interface pocket structure for virtual screening. This approach yielded potent molecule glues that stabilized the CDK12– DDB1 interaction. Together, these findings provide a framework linking endogenous partner identification to interface-guided compound design, potentially expanding the range of recruitable protein pairs for target-directed molecular-glue discovery. Notably, our data outline a versatile and broadly applicable strategy to expand the scope of target protein degradation and thus empower future of chemical induced proximity.

## Results

### Live-cell proximity mapping identifies DDB1 as a candidate transient partner of CDK12

We selected CDK12, a potential therapeutic target in cancer, as a model to investigate whether endogenous transient interactions could provide starting points for molecular-glue design. To identify candidate E3 ligase–associated partners of CDK12, including substrate receptors and adaptor proteins that could contribute to molecular-glue binding^15^, we applied ROProx, an APEX2-based approach that labels proximal proteins in living cells within seconds^13^ (Fig. 1a). Liquid chromatography–mass spectrometry (LC–MS) analysis of affinity-purified labelled proteins from HEK293T cells identified more than 1,000 proteins, including the enriched E3 ligase–associated candidates KEAP1, DDB1 and CBX6 (Fig. 1b). DDB1 enrichment was also observed in HeLa cells (Extended Data Fig. 1a). To prioritize candidates that could bind CDK12 directly and form interfacial pockets suitable for molecular-glue binding, we performed preliminary size-exclusion chromatography (SEC) and cryo-EM analyses. These assessments led us to prioritize DDB1 over KEAP1 and CBX6 (Fig. 2a, b and Extended Data Fig. 2). Subsequent conventional immunoprecipitation–mass spectrometry (IP– MS) showed no co-enrichment of DDB1 with CDK12 under the same condition. The contrast with ROProx is consistent with a transient association and illustrates that live-cell proximity labelling can reveal candidate partners that may be missed by IP–MS (Extended Data Fig. 1b).

**Fig 1:**
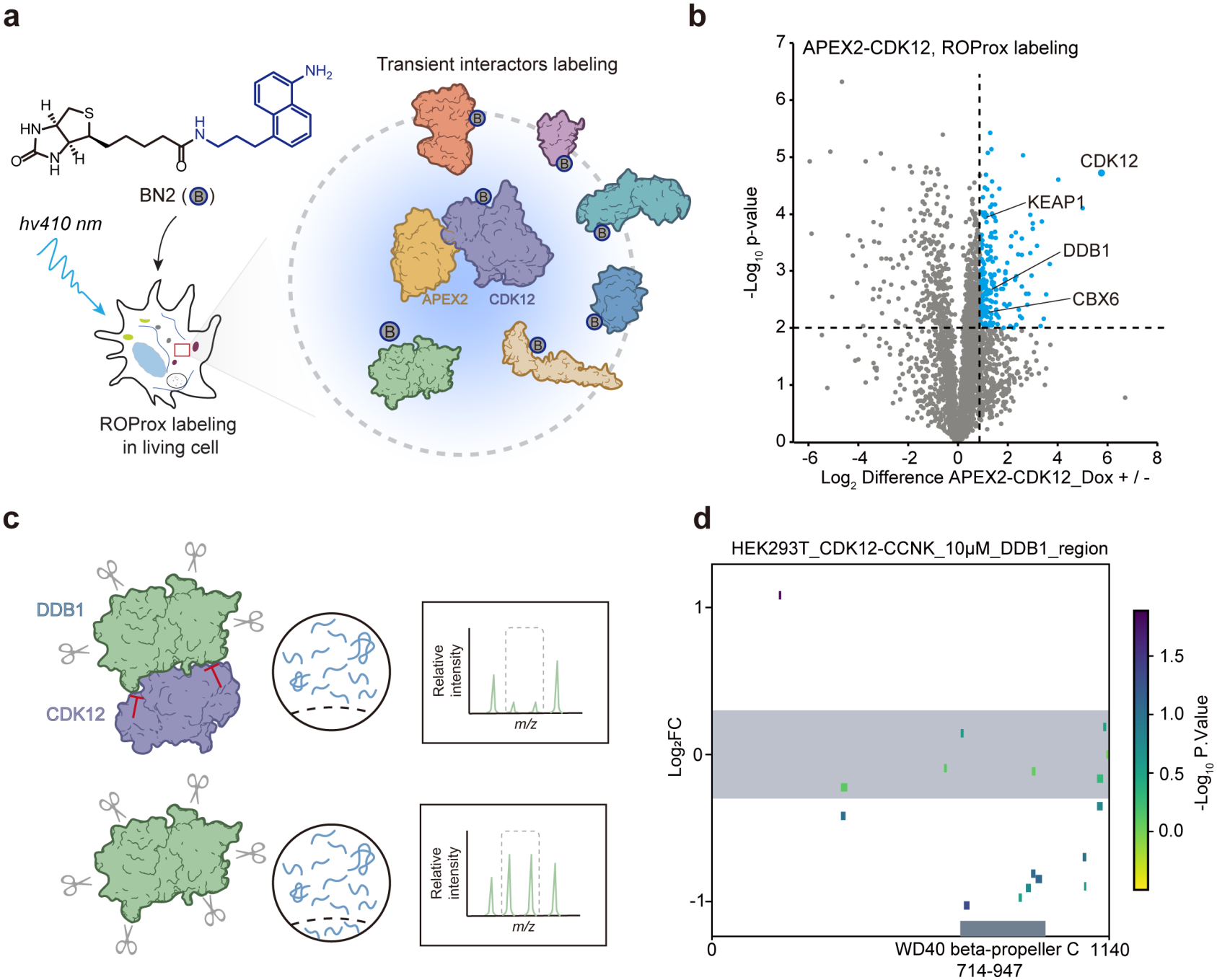
Discovery of DDB1 as a transient CDK12 partnera. **a**, Schematic overview of the ROProx proximity labeling workflow. **b**, Volcano plot showing AP−MS results for the CDK12 interaction labeled by ROProx (biological replicates n = 3, log_2_FC > 1.0, P < 0.05, two-sided Student’s t-test, no adjustments). **c**, Schematic overview of the PELSA workflow. **d**, Local stability profiles to reveal CDK12-binding regions in DDB1. The upper and lower boundaries of the gray-shaded area represent log_2_FCs of 0.3 and −0.3, respectively. The x axis represents the DDB1 sequence from the N-terminal to the C-terminal.

**Fig 2:**
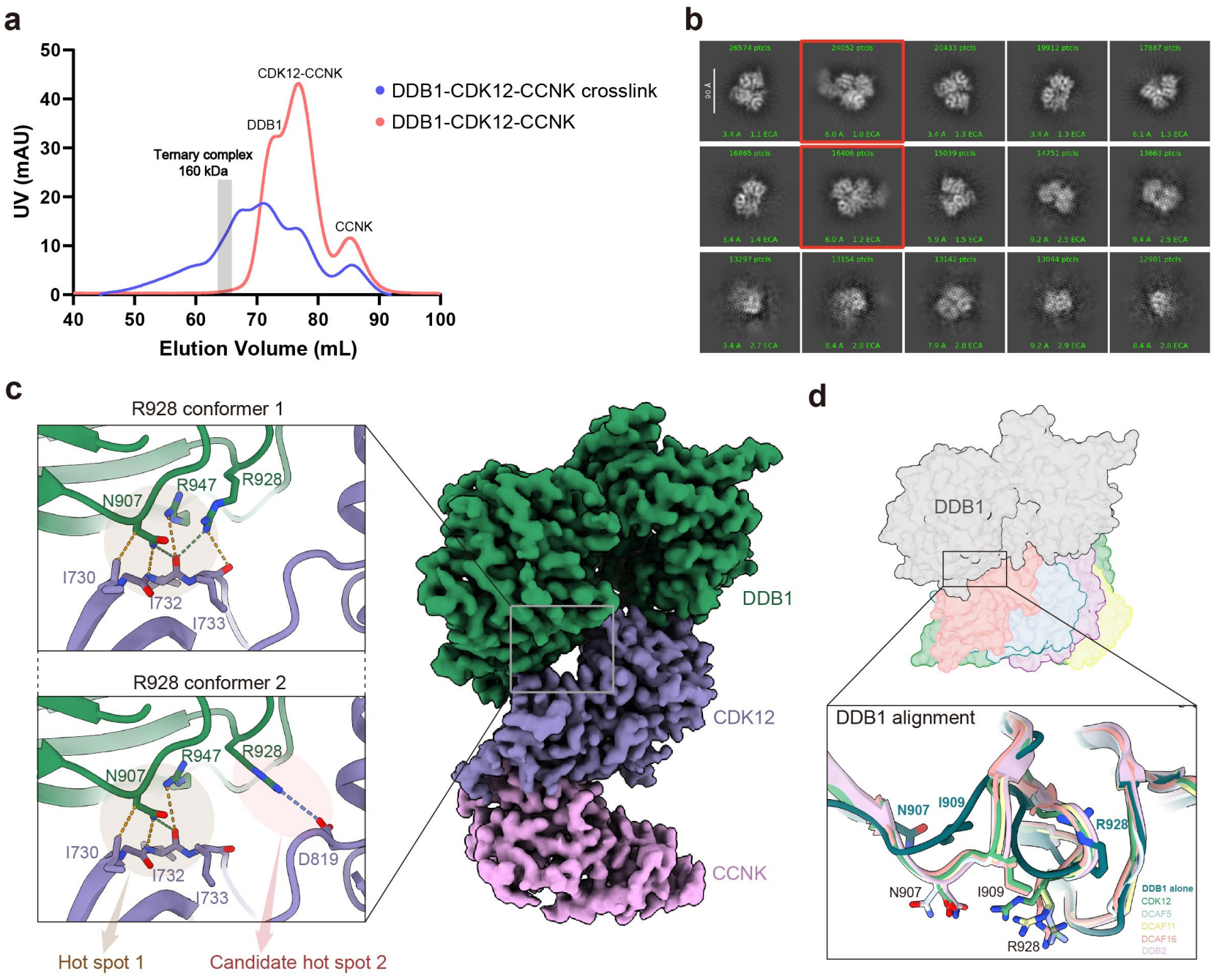
Identification of transient interactions between DDB1 and CDK12. **a**, Size-exclusion chromatography (SEC) profile of DDB1 incubated with the CDK12– CCNK complex with and without BS3 crosslinking. **b**, Top ranked reference-free 2D class averages of the BS3-crosslinked DDB1-CDK12-CCNK complex, with the purified protein sample collected from ternary complex peak in the SEC profile in **a**. The ternary complex class averages are boxed in red. **c**, Left, close-up of the DDB1-CDK12 interface. Hydrogen bonds, electrostatic interaction and other contacts are shown as green, blue and yellow dashed lines, respectively. Right, cryo-EM map of the DDB1-CDK12-CCNK ternary complex. Map obtained from global refinement at a 2.9 Å resolution and sharpened using DeepEMenhancer. **d**, Structure superposition of unoccupied DDB1 (PDB ID: 2B5M) with DDB1 bound to CDK12 and other endogenous substrate receptors (DCAF5: 8TL6, DCAF11: 9PTU, DCAF16: 8G46, DDB2: 3EI1).

To obtain orthogonal biochemical evidence for the CDK12-DDB1 interaction, we performed the peptide-centric local stability assay (PELSA)^14^, which probes local stability shift in proteins upon extensive trypsinization (Fig. 1c). Following incubation of DDB1 with CDK12, PELSA revealed a significant change in proteolytic susceptibility in a region of the DDB1 BPC domain spanning residues 714–739 and 880–947, providing additional biochemical support for the CDK12–DDB1 association (Fig. 1d). Together, these findings support a transient interaction between CDK12 and DDB1 and nominate this protein pair for subsequent structural characterization and interface-guided molecular glue design.

### Transient DDB1-CDK12 contacts prime PPI stabilization

To define the structural basis of the CDK12–DDB1 interaction, we purified DDB1 and the CDK12–cyclin K (CCNK) complex, using the endogenous cyclin partner to stabilize CDK12^16^. Incubation of the purified proteins did not yield a resolvable ternary-complex peak by SEC, whereas BS3 crosslinking enabled collection of the DDB1–CDK12– CCNK complex (Fig. 2a). Together with the ROProx and IP–MS results, these findings support a labile and transient DDB1–CDK12 interaction. Two-dimensional class averages of the crosslinking ternary complex showed a consistent overall architecture (Fig. 2b). Further processing of the cryo-EM data yielded two reconstructions at an overall resolution of 2.9 Å that differed primarily in the side chain conformation of R928 in DDB1, with conformer 1 predominating and conformer 2 accounting for 5% of the classified particles (Fig. 2c).

The structure revealed that CDK12 engages the canonical substrate receptor– binding interface of DDB1^9,15,17-21^, with the ATP-binding pocket of CDK12 positions at and opening toward the protein–protein interface (Fig. 2c). The resolved interface is also consistent with the PELSA results in our study. In the predominant conformer 1, DDB1 N907, R928 and R947 and CDK12 I730, I732 and I733 contribute to a composite interaction hotspot (hotspot 1), with which the side chains of N907 and R928 in DDB1 form hydrogen bonds with the CDK12 backbone (Fig. 2c). TR-FRET measurements also supported the involvement of this interface in CDK12–DDB1 complex formation (Extended Data Fig. 3a). In conformer 2, R928 reorients towards D819 in CDK12, forming a putative electrostatic contact that defines a candidate second hotspot (hotspot 2) and reshapes the interaction network around hotspot 1 (Fig. 2c). This rearrangement highlights the conformational plasticity at the interface and indicates that a molecular glue could bridge or reinforce the reorganized contacts to stabilize the CDK12–DDB1 interaction.

**Fig 3:**
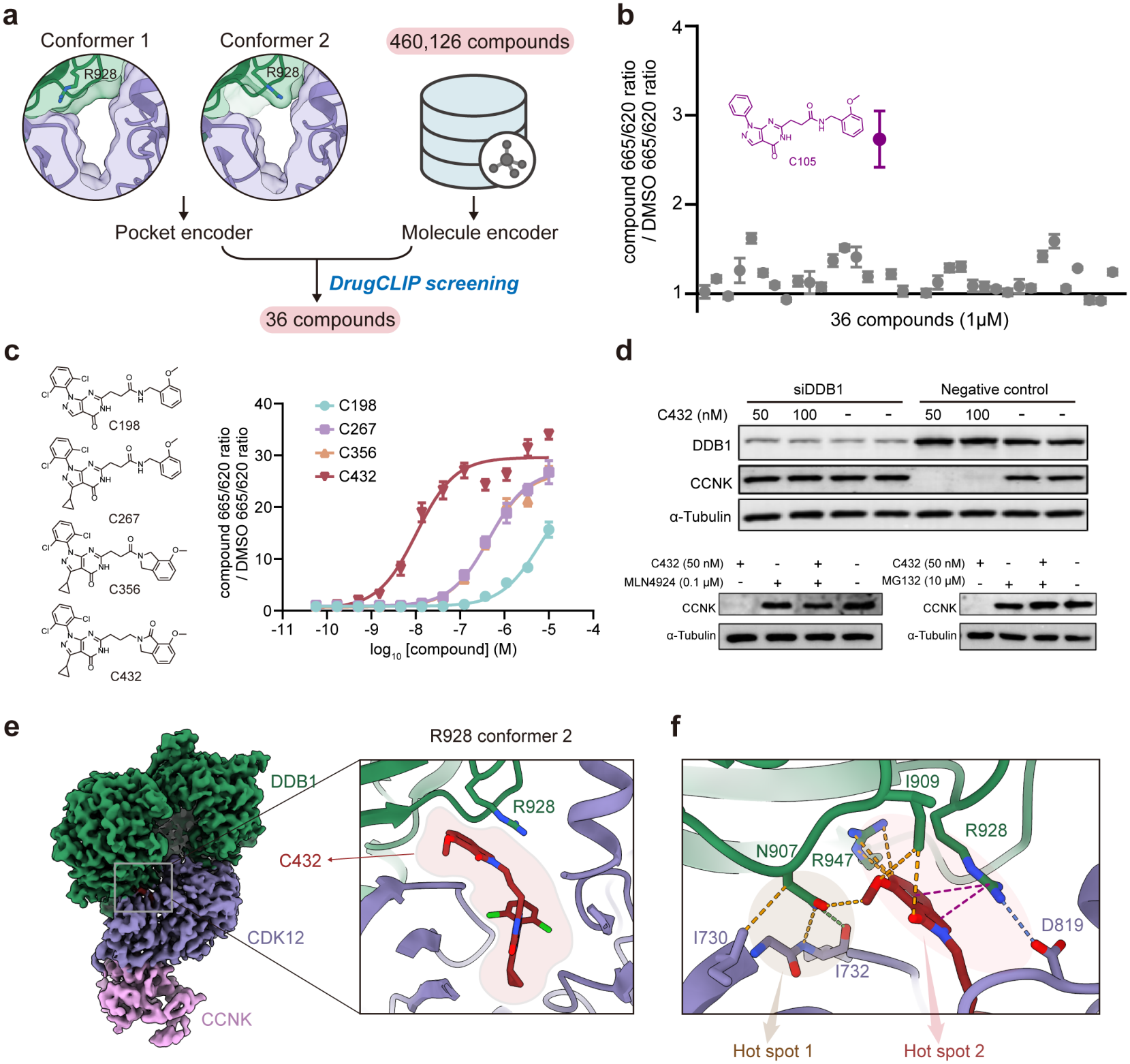
Structure-based molecular gue rational design. **a**, DrugCLIP-based virtual screening of 460,126 compounds using pocket conformations derived from cryo-EM models, yielding 38 compounds for experimental evaluation. **b**, TR-FRET screening of the 38 compounds at 5 μM to assess DDB1-CDK12-CCNK complex formation. **c**, Medicinal chemistry optimization of C105 yield C432, which produced a higher TR-FRET signal than C105. **d**, Immunoblots of CCNK degradation in HCC38 cells treated with or without 50 nM C432, 0.1 μM MLN4924 or 1 μM MG132 for 4 h. Immunoblots of CCNK and DDB1 in HCC38 cells transfected with the DDB1-targeting siRNAs and treated with 50 nM C432. **e**, Cryo-EM structure of the C432-bound DDB1-CDK12-CCNK complex, and close up of the C432 binding mode. **f**, Close-up view of the interaction hotspots at the DDB1-C432-CDK12 interface. Hydrogen bonds, electrostatic interaction, π-cation interactions and other contacts are shown as green, blue, magenta and yellow dashed lines, respectively.

Structural comparison of unoccupied DDB1 with DDB1 bound to endogenous substrate receptors reveal receptor-associated rearrangements of N907, I909, R928 and their surrounding loops (Fig. 2d) ^9,17,19,21,22^. The CDK12-bound DDB1 structure closely resembles the substrate receptor-engaged conformation, suggesting that the transient interaction samples a DDB1 state associated with stable complexes (Fig. 2d). Together, structural characterization of this labile interface revealed pre-existing interaction hotspots and R928 conformational plasticity around the pocket. These features suggest a potential energetic foothold for molecular-glue stabilization of CDK12–DDB1 and support transient endogenous PPIs as rational starting points for glue discovery.

### Transient interface pocket-guided molecular glue rational design

Based on the two cryo-EM models differing in the side-chain conformation of DDB1 R928, we established an ensemble-based virtual screening workflow targeting the CDK12-DDB1 interface pocket (Fig. 3a). Starting from the two pocket models, we used MolCRAFT to generate 10 molecules within each pocket and locally refined the resulting protein-ligand complexes, yielding an ensemble of 20 pocket conformations (Extended Data Fig. 4a). This ensemble was used to screen 460,126 prepared compounds from the Enamine Hit Locator Library using DrugCLIP^23^ (Fig. 3a). Compounds were ranked by their highest normalized DrugCLIP score across the pocket ensemble. Clustering of the top-ranked 1% (4,601 compounds) yielded 86 clusters, from which the two highest-scoring compounds in each of the first 30 clusters were retained (Extended Data Fig. 4b). Subsequent docking-based filtering narrowed this set to 38 compounds (Fig. 3a), which were reranked using AlphaRank to prioritize experimental evaluation.

We next evaluated 38 compounds at 5 μM performing TR-FRET to assess DDB1-CDK12-CCNK complex formation. C105 elicited the largest increases in TR-FRET signal and was selected as the starting point for medicinal chemistry optimization (Fig. 3b). Guided by the proposed binding mode (Extended Data Fig. 4c), we introduced two chlorine substituents on the phenyl ring to restrict its dihedral angle and strengthen the interaction with K756, yielding C198 with improved activity. Further optimization to occupy a small hydrophobic subpocket within the kinase active site afforded C267, followed by cyclization to generate C356 with similar activity. Subsequent modification of the hydrogen-bond donor yielded the optimized compound C432, with an IC_50_ of 22 nM (Fig. 3c). Mechanistic studies further showed that C432 induced CCNK degradation through DDB1-dependent ubiquitination, consistent with previous reports and our design rationale^24,25^ (Fig. 3d).

We next determined a 2.9 Å cryo-EM structure of the C432-bound DDB1-CDK12-CCNK complex, in which DDB1 R928 was resolved exclusively in conformer 2, creating space to accommodate C432 at the interface (Fig. 3e). At the previously proposed candidate hotspot 2, DDB1 N907, I909, R928 and R947 contact with C432, while R928 forms an electrostatic contact with CDK12 D819 (Fig. 3f). Together with hotspot 1, this C432-reinforced interaction network provides a structural basis for stabilization of the CDK12–DDB1 interaction, as also supported by TR-FRET measurements (Extended Data Fig. 3b).

## Discussion

Transient protein–protein interactions are prevalent, yet effective ways to regulate and exploit them remain limited. Molecular glues are small molecules that can stabilize these interactions, thereby promoting target protein degradation or modulating protein function. Previous molecular glues discovery has largely followed compound-first approaches, in which known compounds or compound libraries are evaluated through phenotypic, proteomic or affinity-based screening before the suitable POI-E3 pairs are identified. In this study, we present TrIP–MGRD, a framework that identifies endogenous transient PPI as starting points for molecular-glue discovery. MS–based proteomics identified DDB1 as an E3 ligase–associated partner of CDK12. The complex structure revealed their direct contacts that may provide an energetic basis for PPI stabilization, as well as an interfacial pocket suitable for virtual screening. DrugCLIP screening and structure-based optimization yielded a potent molecular glue C432 efficiently.

MS-based proteomic approaches can identify interaction partners and candidate neo-substrates, providing detailed information for molecular-glue discovery^26^. This depth of information, however, often comes at the expense of chemical throughput. By contrast, fluorescence-based interaction assays can support high-throughput compound screening, but assay development generally requires a predefined protein pair and a suitable interaction readout^11^. Similarly, interface-guided virtual screening can evaluate large compound libraries but depends on a suitable protein pair and a structural model of its interface^23^. These complementary strengths support a sequential strategy in which proteomic profiling first identifies candidate endogenous partners of a target. Characterization of these interactions and their interfaces provides an early basis for assessing the target’s potential for molecular-glue modulation and prioritizing suitable protein pairs before extensive compound screening. Large chemical libraries can then be screened against the selected pairs using fluorescence-based interaction assays or interface-guided virtual screening. By focusing the chemical search on defined interactions, this approach could improve discovery efficiency and make compound screening more scalable.

Characterizing the pocket at a transient protein–protein interface helps assess whether it can accommodate a molecular glue that stabilizes the interaction. Our findings reveal the contacts underlying the CDK12–DDB1 association and identify interfacial features that can guide the design of such compounds. In the compound-free DDB1–CDK12–CCNK structure, DDB1 residues form direct contacts with CDK12 that resemble those observed in stable protein complexes^9,17,19,21,22^. TR-FRET measurements support the contribution of this interaction region, which we designate hotspot 1. An alternative R928 orientation defines a second potential interaction region (candidate hotspot 2). In the glue-bound structure, the R928 side chain adopts the less populated conformation observed in the compound-free complex. Contacts involving R928 and the bound glue reinforce hotspot 2 alongside the pre-existing hotspot 1, forming the proposed “twin hotspots” arrangement. These observations support a model in which transient protein–protein contacts provide a potential energetic foothold for stabilization, while molecular-glue binding contributes a complementary interaction hotspot that strengthens the PPI.

Molecular glues hold considerable therapeutic promise, yet their rational design remains challenging. Our findings support the use of transient protein–protein interactions as starting points for rational molecular-glue design. The CDK12–DDB1 example shows how characterization of a weak interface can guide the discovery of compounds that strengthen the association. Extending TrIP–MGRD to other E3 ligase– POI pairs could help identify additional interfaces amenable to chemical stabilization and broaden the range of interactions explored for molecular-glue discovery.

## Acknowledgements

We thank the staff members of the Electron Microscopy System (https://cstr.cn/31129.02.NFPS.EMAS) at the National Facility for Protein Science in Shanghai (https://cstr.cn/31129.02.NFPS), for providing technical support and assistance in data collection and analysis of apo and C432 induced DDB1-CDK12-CCNK complexes. We acknowledge the financial support from the National Natural Science Foundation of China (82530108, 22521104 to K.D., 22534003, 22541703 to R.T.), the National Key R&D Program of China (2023YFE0119000, 2023YFF1205104, 2023YFC2506402 to K.D., 2024YFA1307204 to R.T.), the Strategic Priority Research Program of the Chinese Academy of Sciences (XDB1060000 to K.D.), the State Key Laboratory of Chemical Biology, Innovative Drug Research and Development-National Science and Technology Major Project (2025ZD1802501 to Y.L.), Beijing Frontier Research Center for Biological Structure Fundings, Postdoctoral Fellowship Program of CPSF (GZB20260867 to S.J.), the innovation program of science and research from the DICP, CAS (DICP I202521 to X.Z.).

## Author contributions

K.D. conceived the project, with the collaboration of R.T. for characterizing transient interaction via ROProx method, Y.L. for leading the application of the DrugCLIP method to virtual screening and M.Y. for characterizing interaction region of the CDK12-DDB1 complex via PELSA method. K.D., R.T., Y.L., M.Y. and Z.L. managed and designed the experiments. Z.L., S.J., M.K., W.Z. and X.Z. performed most of the experiments. S.J., J.Z. and G.W. designed the constructs and purified these protein complexes. S.J. performed cryo-EM data processing and model building. M.K. and P.Z. performed H2O2-independent ROProx proximity labeling experiment. Y.L. and X.S. performed mechanism study. G.W. performed TR-FRET experiment. Y.Y. performed PELSA experiment and statistical analyses. Z.L. conducted the compounds optimization. Z.L., Y.C. and Y.C. performed chemical synthesis. W.Z. and Z.L. led the application of the DrugCLIP method to virtual screening. M.T. and W.Z. supported the MS study. K.D., Z.L. and S.J. wrote the manuscript.

